# Bone pain in Fibrous dysplasia does not rely on aberrant sensory nerve sprouting or neuroma formation

**DOI:** 10.1101/2024.07.18.603554

**Authors:** Biagio Palmisano, Chiara Tavanti, Giorgia Farinacci, Giorgio Gosti, Marco Leonetti, Samantha Donsante, Giuseppe Giannicola, Natasha Appelman-Dijkstra, Alessandro Corsi, Ernesto Ippolito, Mara Riminucci

**Author notes:** Tettamanti Center, Fondazione IRCCS San Gerardo dei Tintori, 20900 Monza, Italy (Samantha Donsante).

## Abstract

Bone pain is a major symptom of many skeletal disorders. Fibrous dysplasia (FD) is a genetic disease with mono or polyostotic skeletal phenotype due to the post-zygotic occurrence of the causative Gsα mutation. Bone pain in FD often associates with skeletal deformities and fractures or nerve impingement by the pathological tissue. However, even in the absence of complications, FD patients often complain of a chronic pain that does not correlate with their disease burden. Multiple hypotheses have been made to explain this pain. However, its pathogenetic mechanisms remain, as yet, largely unexplored.

In this study, we first demonstrate that the FD mouse model EF1α-Gsα^R201C^ develops a painful-like behavior and an altered response to nociceptive stimuli that, as in FD patients, do not correlate with the severity of their phenotype, thus providing a reliable model to study bone pain in FD. Then, we show that in EF1α-Gsα^R201C^ mice, the overall pattern of skeletal innervation is preserved and that within FD lesions, sensory fibers are variably and focally distributed, mainly at perivascular sites. Finally, we provide the first analysis of a series of human FD bone biopsies showing that sensory nerve fibers are rarely detected within the pathological tissue.

These data confirm that bone pain is an intrinsic and reproducible feature of FD. They also show that, albeit sensory nerve fibers are found within FD lesions and may contribute to the unpleasant sensation that accompanies the disease, pathological sensory nerve sprouting or formation of neuromas are not detected in the Gsα-mutated skeleton.

## Introduction

Bone pain is a common and severe symptom of many bone diseases, from genetic disorders to cancer metastasis. As in other organs, the sensation of pain in bone mainly results from signals that are collected by the peripheral nervous system upon stimulation (nociceptive pain) or damage (neuropathic pain) of different types of nerve fibers and are then processed by the central nervous system. Studies in humans^1^ and animal models^2–4^ showed that bone pain is conducted mostly by A-delta and C nerve fibers. The former are myelinated fibers that express the nerve growth factor (NGF) receptor tropomyosin kinase A (TRKA) and the 200 kDa neurofilament (NF200); the latter are unmyelinated fibers that express TRKA and calcitonin gene related peptide (CGRP)^4,5^. Both A-delta and C fibers are found with decreasing density in the periosteum, bone marrow and cortical bone. In the periosteum they can detect mechanical stimuli, e.g. in bone fractures, while in the bone marrow they conduct sensory information due to compression, e.g. by cancer cells. In some pathological conditions, bone nerve fibers may undergo sprouting processes or may form neuroma-like structures that further increase the sensation of pain.

Bone pain is a major clinical aspect of Fibrous dysplasia (FD) of bone, a rare, skeletal disease caused by gain-of-function mutations of the Gsα gene, most commonly at the codon R201 (R201C, R201H)^6^. Due to the post-zygotic occurrence of the mutations, FD patients are clinically heterogeneous with different mono or polyostotic patterns of skeletal involvement and, in the most severe cases, different types of extra-skeletal manifestations (McCune-Albright syndrome, MAS)^7^. The development of FD lesions occurs through resorption of normal bone and depletion of hematopoietic cells and marrow adipocytes that are all replaced by fibrous tissue containing tiny and hypo-mineralized bone trabeculae. A previous report showed that both adults and children with FD/MAS may experience pain^8^. In addition, an analysis performed on a large cohort of adult FD patients, demonstrated the presence of both nociceptive and neuropathic pain, respectively in 45,4% and 31,3% of the participants^9^ while another study reported that pain in FD/MAS may show neurobiological properties and neuropsychological features^10^. Despite the presence of pain and its variable and often unsatisfactory response to pharmacological treatments such as anti-resorptive, anti-inflammatory drugs and opiates^11–14^, the mechanisms of pain sensation in FD patients are, as yet, incompletely understood. Furthermore, although it has been assumed that the pathological fibro-osseous tissue associates with abnormal nerve fiber growth^11^, no data are currently available on the pattern of innervation of FD lesions. Transgenic mouse models may then represent a valid strategy to explore the characteristics and morphological bases of bone pain in this rare condition.

The EF1α-Gsα^R201C^ mouse model reproduces the elementary tissue changes of human FD, shows time-dependent heterogeneous patterns of skeletal involvement as FD patients, and has been proved to be a useful tool for investigating the effect of pharmacological strategies on the development and progression of the disease^15–17^.

In the present work, we report that EF1α-Gsα^R201C^ mice develop a pain-like behavior and an altered nociceptive response that, as in humans^18^, does not correlate with their skeletal disease burden. Moreover, by using the new *BAF53b-GFP;EF1α-Gsα^R201C^* reporter mouse model with GFP-expression in nerves along with immunolocalization studies, we show that the pattern of sensory innervation of skeletal compartments such as periosteum and bone marrow is overall normal in the Gsα mutated mice whereas within FD lesions, nerve structures, including sensory fibers, are focally detected along arterial branches. Finally, we show for the first time that in human FD lesions, sensory innervation is limited and mainly associated with large arterial vessels.

## Methods

### Generation of experimental mice

Generation and characterization of the EF1α-Gsα^R201C^ mouse model were reported previously (Saggio et al JBMR 2014). *Tg(Actl6b-Cre)4092Jiwu/J (BAF53b-Cre)* mice, a transgenic line expressing Cre recombinase under the control of the pan-neuronal *Actl6b* gene promoter (*BAF53b-Cre*)^19^, and the reporter line *B6.129(Cg)-Gt(ROSA)26Sortm4(ACTB-tdTomato,-EGFP)Luo/J* (*R26-mTmG)*^20^ were purchased from The Jackson Laboratory (Bar Harbor, Main, USA). *BAF53b-Cre mice* were crossed with the *R26-mTmG mice* reporter line and then with *EF1α-Gsα^R201C^* mice to generate *BAF53b-GFP;EF1α-Gsα^R201C^* mice, a triple transgenic mouse model of FD expressing GFP in all nerve fibers. These mice were maintained at a mixed background (B6;FVB), therefore only littermates were used as controls. Genotyping of mice was performed on 3-week-old mouse tail tip by PCR analysis using MyTaq DNA^TM^ Polymerase (Meridian Bioscience Inc., Cincinnati, Ohio, USA) and specific oligonucleotide primers (Table S1).

All animals were maintained in cabin-type isolators at standard environmental conditions (temperature 22–25°C, humidity 40–70%) with 12:12 dark/light photoperiod. Food and water were provided *ad libitum*. All studies were performed in compliance with relevant Italian laws and Institutional guidelines and all procedures were IACUC approved.

### Behavioral tests

All behavioral analyses were carried out during the dark phase, starting from 6 pm, to avoid any potential behavioral inhibition of the resting light phase^21^. Experimental groups were generated by random selection of transgenic mice, to allow the inclusion of EF1α-Gsα^R201C^ mice with diverse skeletal phenotypes and avoid potential biases. The number of mice used for each test are reported in the figure legends.

#### Burrowing test

This test assays the spontaneous ability of mice to empty a tube filled with food pellets. The test was performed as previously described^22^ with minor modifications. To generate the burrow tubes, a PVC downpipe with 68 mm diameter (OBI, Rome, Italy) was cut into 20 cm long lengths. One end of each tube was sealed with an aluminum lid and the other end was elevated about 3 cm off the floor with 5-cm long machine screws. Habituation was carried out O/N and was conducted in group by placing 4-5 mice into a 26 (length) × 23 (width) × 17 (height) cm cage containing the burrow tube filled with food pellets. Forty-eight hours later each mouse was transferred into a cage containing the tube filled with 300 grams of pellets. The evaluation of the burrowing activity was performed by weighing the food remaining in the tube at different time points. The first evaluation was performed after 2 hours (2H), then the tube was filled again with pellets and a second measurement was taken the next morning (ON).

#### Nesting test

This test evaluates the ability of mice to build a nest. Mice were transferred into individual cages containing 4,5 grams of orderly placed shredded paper as nesting material without any other environmental enrichment. The following morning, the nest was photographed and given a score ranging from 1 to 5^23^ based on the organization and the amount of paper used to build it. A detailed table is reported in Fig. S1A.

It must be considered that nest construction reflects a complex interaction between the animal and its environment and in some cases the results did not fit perfectly one of the scores. In these cases, a half point was either subtracted or added to the score^23^. For example, some mice built a score 5 nest but used a bit less than 90% of the paper thus receiving a score of 4.5.

#### Open field test

This test allows to evaluate the overall motor activity of mice. Mice were transferred to the testing room at short distance from the holding facility. Each mouse was placed into the center of a 26 (length) × 15,5 (width) × 14 (height) cm cage and allowed to explore it freely for 5 minutes for habituation. Then, it was monitored for 2 minutes by video tracking. Videos were filmed using a high-end single-cam device, featuring a 12Mp 1/2.55″ sensor (mounted on an Apple iPhone XR), at 1920 × 1080 resolution, 30 frames/second and analyzed by Python code developed in house using Numpy, Matplotlib, Pandas, and OpenCV. First, the mouse body was segmented using the HSV color space. We then computed the mouse center and the distance traveled by the mouse and tracked the time spent close to the cage walls as opposed to the center. The code is available online at https://github.com/ggosti/HueMouse. The overall motor activity was quantified as the total traveled distance.

#### Tail flick test

This test measures the latency of the avoidance response, i.e. the flicking of the tail when pain is induced by a radiant heat. Each mouse was gently held and positioned on the tail flick apparatus (Ugo Basile, Gemonio, Italy) with the tail placed over a small window emitting a beam of I.R. energy at 65% intensity. The heat stimulus was applied on the mid portion and on the tip of the tail and the latency of the tail flicking was automatically scored by an optic fiber.

### X-ray analyses and disease burden score

The disease burden score was assessed on radiographic images by an arbitrary scoring system. Images were taken by Faxitron MX-20 Specimen Radiography System (Faxitron X-ray Corp., Wheeling, IL, USA) set at 24-25 kV for 6-8 seconds under anesthesia with a mixture of Zoletil 50/50 (Virbac SA, Carros, France) and Rompun (Bayer, Leverkusen, Germany). A value from 0 to 3 was assigned to each bone segment based on the area occupied by FD lesions as reported in Fig. S2. Specifically, score 0 was assigned when no FD lesions were detected; score 1 when lesions involved less than 25% of the bone segment; score 2 when FD lesions occupied 25-50% and score 3 when more than 50% of the bone segment was affected. All skeletal segments were analyzed and the sum of individual bone scores was considered as the final score for each mouse.

### Histology and immunofluorescence of mouse samples

Mice were euthanized by carbon dioxide inhalation and the skeletal segments were fixed with 4% formaldehyde solution for 6 hours at 4°C. After decalcification in 0.5M EDTA for 48-96 hours, the samples were embedded in porcine gelatin as described previously^24^ and 60-μm-thick sections were cut and kept at -20°C. To visualize the endogenous fluorescence, sections were thawed at RT for 30 minutes, rehydrated with PBS for further 30 minutes and stained for 30 minutes with the nucleic acid stain TO-PRO3 (T3605, Thermo Fisher Scientific, Waltham, MA, USA) for nuclei visualization.

For immunofluorescence, sections were thawed for 20 min, rehydrated in PBS with 0.3% Triton for 10 minutes and incubated with 5% goat serum for 30 minutes at RT. The anti-NF200 antibody (N4142, Sigma-Aldrich, Saint Louis, MO, USA) was applied at a dilution of 1:500, the anti-TRKA antibody (AF1056, R&D System, Minneapolis, MN, USA) was diluted 1:50 whereas for the anti-CGRP antibody (ab81887, Abcam, Cambridge, UK) a dilution of 1:1000 was used. Anti-SCA-1 and anti-CD31 antibodies (122501 and 102501, Biolegend, San Diego, CA, USA) were applied at a dilution of 1:100. Incubation of all primary antibodies was carried out O/N at 4°C. Sections were then washed several times with PBS, incubated at RT for 1.5 hours with appropriate secondary antibodies (A-11034 and A-1178, Thermo Fisher Scientific), washed again with PBS and stained with TO-PRO-3 (T3605, Thermo Fisher Scientific) for 20 minutes to visualize nuclei. All images were acquired with Leica Confocal Microscope (Leica Biosystems, Wetzlar, Germany). Z-stacked images were analyzed with Fiji ImageJ software^25^. The analysis of nerves and blood vessels was performed by randomly selecting areas in caudal vertebrae from BAF53b-GFP and *BAF53b-GFP;EF1α-Gsα^R201C^* mice. Nerve fibers and blood vessels were manually traced to assess the ratio between their respective lengths.

### Human samples

FD and control bone samples were used. The former were the same archival paraffin embedded samples used in our previous study^26^. The latter were collected from healthy donors and obtained as surgical waste during different types of orthopedic surgery. Samples were used following patients’ written informed consent and with the approval of the local Institutional Review Board.

### Histology and immunohistochemistry of human FD lesions

Histological studies on human FD included 13 FD bone biopsies collected from femurs and tibiae of 7 FD/MAS patients, 5 males and 2 females, age range 6-23 years, some of them receiving bisphosphonates at the time of biopsy collection. Patients complained of pain at affected sites at the time of surgery, although no assessment through standard evaluation scales was performed.

After fixation with 4% formaldehyde at 4°C and decalcification in 0.5 M EDTA, biopsies were embedded in paraffin according to standard procedure. One sample was embedded in porcine gelatin according to the same procedure used for mouse tissues. Histological analysis was performed on 3 μm-thick sections cut from paraffin blocks after Hematoxylin-Eosin and Sirius red staining.

Immunohistochemistry was performed on paraffin sections after heat-induced epitope retrieval in citrate buffer. For NF200 staining, sections were incubated with anti-NF200 antibody (NCL-L-NF200-N52, Leica Biosystems) applied at a dilution of 1:50 for 30 minutes at 25 °C in an automated Leica BOND III system (Leica Biosystems). Visualization of the primary antibody binding to tissue section was performed using BOND Polymer Refine Detection (DS9800, Leica Biosystems). For CGRP immunolocalization, a mouse anti-human antibody (ab81887, Abcam) was applied at the dilution of 1:200 for 2 hours at room temperature. After incubation with the primary antibody and repeated washing with PBS, sections were exposed for 30 min to biotin-conjugated polyclonal rabbit anti-mouse (E0464, Agilent Dako) 1:300 in PBS and then for 30 min to horseradish peroxidase-conjugated streptavidin (P0397, Agilent Dako) 1:1000 in PBS. The HRP reaction was developed using 3,3’-diaminobenzidine tetrahydrochloride kit (SK-4105, Vector Laboratories, Burlingame, CA, USA) as chromogen.

Immunolocalization on 60 μm-thick gelatin sections was performed using anti-NF200 antibody (N4142, Sigma-Aldrich) following the same protocol described for mouse sections.

### Gene expression analysis

Human bone specimens were homogenized by Mikro-Dismembrator U (Sartorius, Gottingen, Germany) and total RNA was isolated using the TRI Reagent (ThermoFisher Scientific) protocol. To obtain cDNA, reverse transcription was performed using PrimeScript RT Reagent Kit (Takara) according to manifacturer’s protocol. Gene expression was determined by quantitative PCR (qPCR) analysis on a 7500 Fast Real-Time PCR System (Applied Biosystem), using PowerUP Sybr Green (Thermo Fisher Scientific) and specific primers (Table S1). Gene expression levels of each gene were normalized to RNA 18s ribosomal N5 (*RNA18SN5*) expression.

### Statistical analysis

Non-parametric Mann Whitney test was used to compare two groups when the population of data did not have a normal distribution while unpaired Student’s t-test was used when the two populations were normally distributed. Two-way ANOVA test was used to detect statistical differences in the behavioral tests between WT and EF1α-Gsα^R201C^ mice at different ages. Correlation analyses were conducted using the Pearson test and coefficient r and p-value were showed. In all experiments a p-value <0.05 was considered statistically significant. All graphs and statistical analyses were performed using GraphPad Prism version 10 (GraphPad Software, La Jolla, CA, USA).

## Results

### EF1α-Gsα^R201C^ mice develop spontaneous bone pain-like behavior

The development of pain-like behavior in a mouse model of FD (EF1α-Gsα^R201C^ mouse) was assessed using standard tests. Burrowing capacity resulted significantly reduced in EF1α-Gsα^R201C^ compared to WT mice, in both males and females (Fig. 1A-D). Age stratification of mice revealed that burrowing behavior in EF1α-Gsα^R201C^ female mice was reduced as early as 3 months of age (Fig. 1B), whereas differences in males were observed starting at 5 months of age (Fig. 1D). Two-way ANOVA simple main effects analysis indicated that genotype, rather than age, had a statistically significant effect on the burrowing capacity in both female and male mice (Fig. 1B, D).

**Figure 1.**
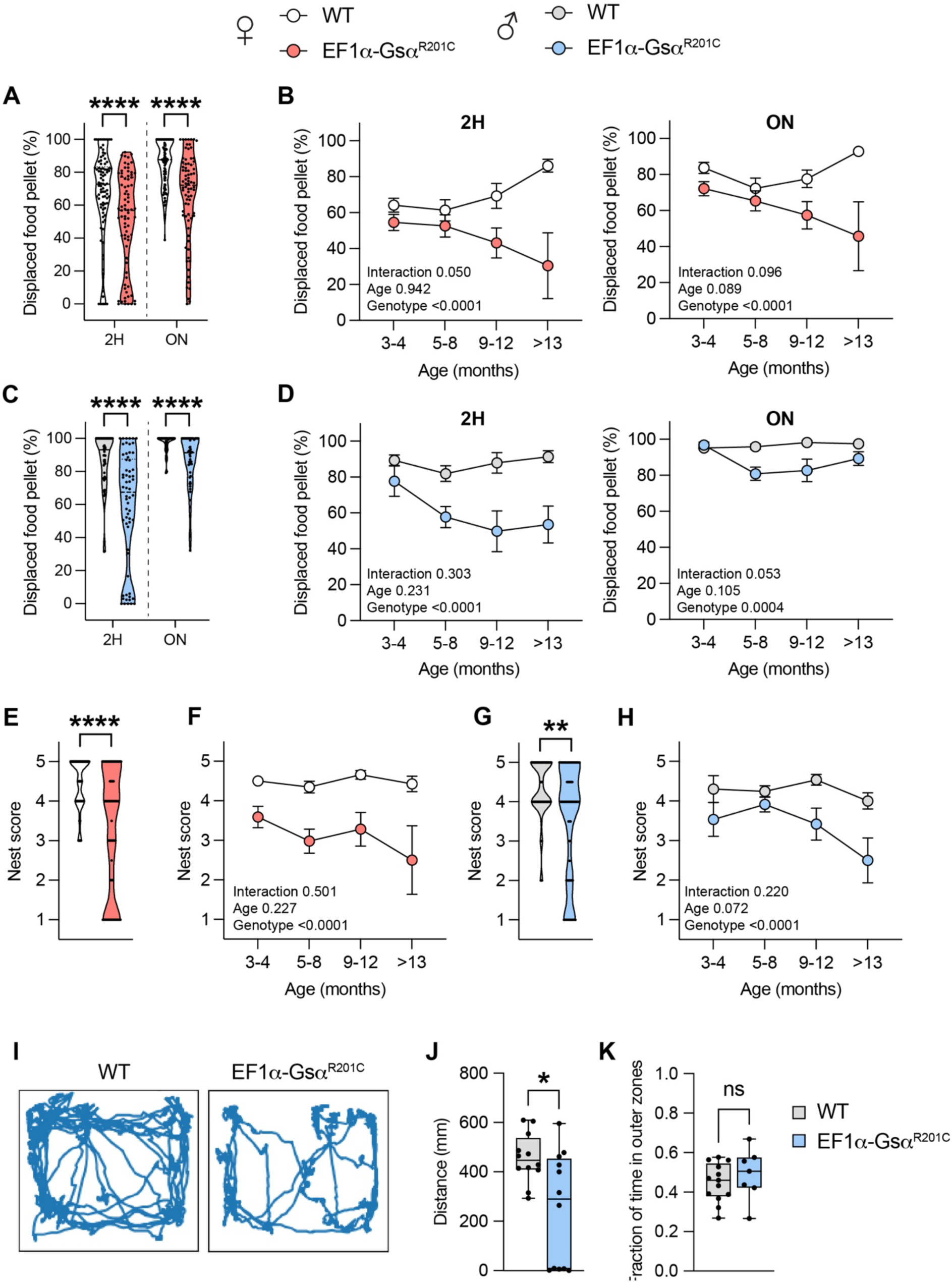
Assessment of spontaneous pain-like behavior. A) Burrowing test results obtained from 2-hour (2H) and overnight (ON) experiments on female mice. WT n=90; EF1α-Gsα^R201C^ n=78, Mann-Whitney test ****p<0.0001. B) Age stratification of burrowing test results from female mice. P-values from the two-way ANOVA analysis are reported in each graph. C) Burrowing test results obtained from 2H and ON experiments on male mice. WT n=56; EF1α-Gsα^R201C^ n=62, Mann-Whitney test ****p<0.0001. D) Age stratification of burrowing test results from male mice. P-values from the two-way ANOVA analysis are reported in each graph. E) Nesting test results from experiments performed on female mice. WT n=83; EF1α-Gsα^R201C^ n=77, Mann-Whitney test ****p<0.0001. F) Age stratification of nesting test results from female mice. P-values from the two-way ANOVA analysis are reported in each graph. G) Nesting test results from experiments performed on male mice. WT n=67; EF1α-Gsα^R201C^ n=70, Mann-Whitney test ****p<0.0001. H) Age stratification of nesting test results from male mice. P-values from the two-way ANOVA analysis are reported in each graph. I) Representative video tracking images of OFT performed in male mice. J) Total distances covered by the mice during the 2-minutes OFT. WT n=12; EF1α-Gsα^R201C^ n=12, Mann-Whitney test *p<0.05. K) Fraction of time spent by the mice in the outer zones of the cage. For this calculation, immobile mice were excluded from the analysis.

Similarly, the capacity of both female and male EF1α-Gsα^R201C^ mice to build a well-formed nest was significantly compromised (Fig. 1E-H). Nests built by WT mice were three-dimensional, well assembled and developed in height (Fig. S1B). In contrast, most of the nests made by EF1α-Gsα^R201C^ mice were flat and disorganized and in some of them the paper strings were chewed by the mice (Fig. S1B). Stratification of results according to age demonstrated lower nesting score in EF1α-Gsα^R201C^ mice compared to WT at all ages and in both sexes (Fig. 1F, H). Two-way ANOVA simple main effects analysis showed that genotype, but not age, had a statistically significant effect on this behavioral test, in both female and male mice (Fig. 1F, H). Furthermore, the analysis of mouse ambulation performed by open field test indicated that EF1α-Gsα^R201C^ mice had an impaired locomotor activity compared to WT (Fig. 1I, J). However, the fraction of time spent in the outer zones of the cage was similar between the two genotypes thus indicating the absence of an anxious-related behavior in transgenic mice^27,28^ (Fig. 1K). Altogether, these data demonstrated that EF1α-Gsα^R201C^ mice developed a spontaneous pain-like behavior.

### EF1α-Gsα^R201C^ mice show delayed nociceptive response

In order to assess if FD lesion development in mice altered their nociceptive response, we used the tail flick test. The test was performed on two segments of the mouse tail, the mid-tail and the tail tip, and the results were analyzed independently. Interestingly, we observed an overall longer latency of response to the heat stimulus in EF1α-Gsα^R201C^ compared to WT mice (Fig. 2A-D). This result was more consistent in males in which the reaction time was significantly increased compared to WT at both mid-tail and tail tip (Fig. 2C), while in females a statistically significant difference was observed only at the tail tip (Fig. 2A). Two-way ANOVA analysis revealed that, except for the mid-tail measures in females, genotype had a statistically significant effect on the tail flick test (Fig. 1B, D).

**Figure 2.**
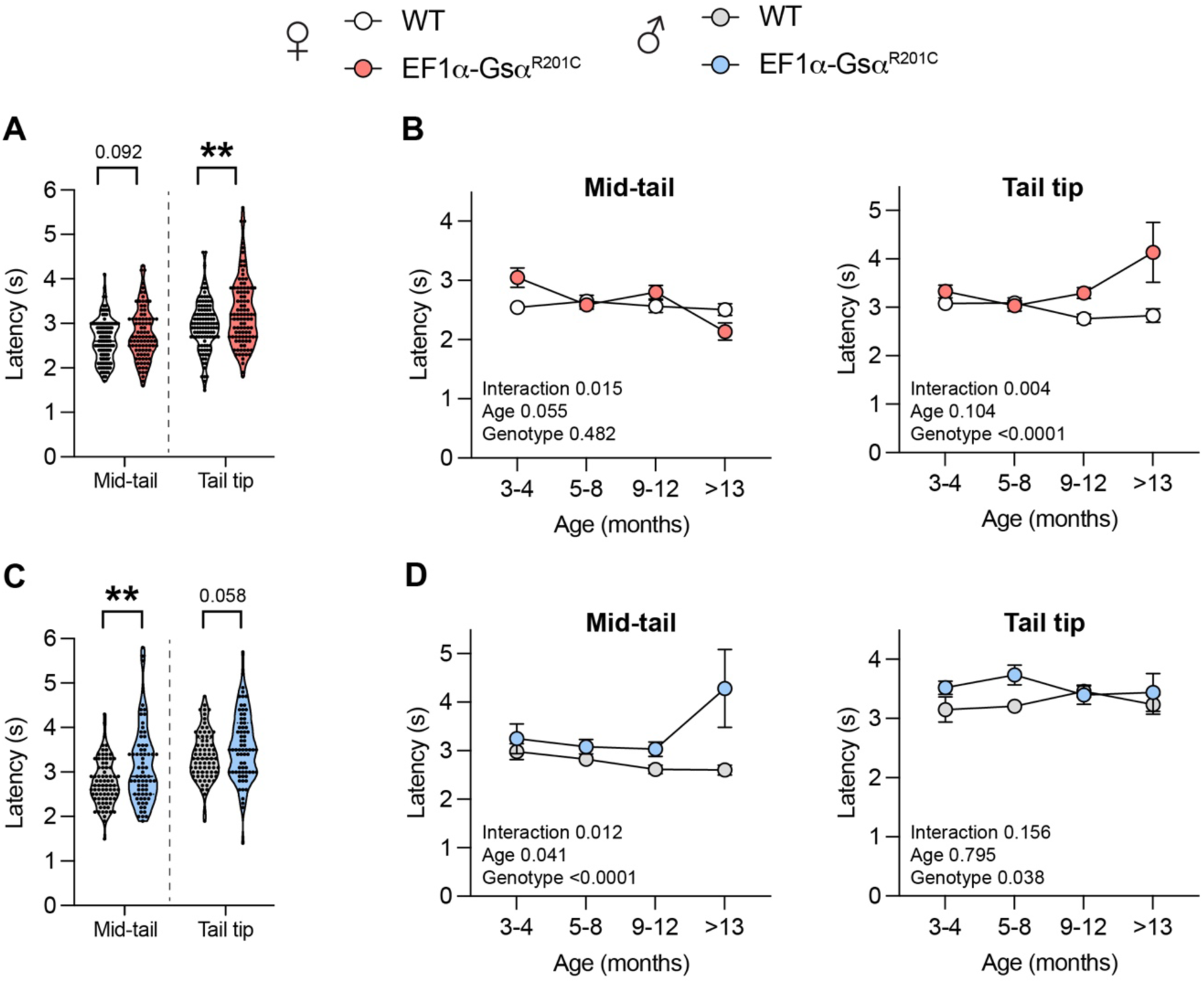
Assessment of nociceptive response. A) Tail flick test results obtained on the mid-tail and tail tip of female mice. WT n=110; EF1α-Gsα^R201C^ n=99; Welch’s t test **p<0.01 or exact p-value is reported. B) Age stratification of tail flick test results from female mice. P-values from the two-way ANOVA analysis are reported in each graph. C) Tail flick test results obtained on the mid-tail and tail tip of male mice. WT n=80; EF1α-Gsα^R201C^ n=71; Welch’s t test **p<0.01 or exact p-value is reported. D) Age stratification of tail flick test results from male mice. P-values from the two-way ANOVA analysis are reported in each graph.

These data demonstrate that EF1α-Gsα^R201C^ mice did not show increase nociception and rather displayed a delayed response to stimulation of sensory fibers.

### Bone pain-like behavior in EF1α-Gsα^R201C^ mice is not correlated with FD disease burden

To assess the correlation between the skeletal disease burden of EF1α-Gsα^R201C^ mice and their age, body weight and pain-like behavior we developed a disease scoring system based on radiographic analysis (Fig. S2A). Pearson correlation coefficient revealed a significant, positive correlation between the skeletal burden score and the mouse age (Fig. S2B). This confirmed the validity of the disease scoring method as the number and severity of FD lesions in EF1α-Gsα^R201C^ mice increase with age^29^. Accordingly, a positive although not statistically significant correlation was also observed between skeletal burden and body weight (Fig. S2C), as the latter tends to increase with age in rodents. This finding also indicates that albeit EF1α-Gsα^R201C^ mice are markedly lighter than WT (Fig. S2D), they retain the ability to gain weight during ageing in spite of the progression of the disease (Fig. S2D).

Then we analyzed the correlation between the results of the behavioral tests and the skeletal disease burden scores. Surprisingly, no correlation was found between the spontaneous pain-like behavior, as assessed by the burrowing and nesting tests, and the extent of skeletal involvement by the disease (Fig. S2E, F). Interestingly, we observed that mice showing the same disease burden scores could present variable burrowing or nesting capacity. For instance, mice with lesions detected only in tail could show the worst pain-like behavior, while mice with many skeletal segments affected and high degree of disease burden could experience no pain at all (Fig. S2E, F). As in the other tests, no significant correlation was observed between the skeletal disease burden score and the results of the tail flick test (Fig. S2G).

### Nerve fibers distribution within FD bone lesions

To investigate whether the pattern of innervation of mouse bones was altered in Gsα-mutated mice, either as a consequence of FD lesion development or as a cell autonomous effect of the mutation in the peripheral nervous system, we set up lineage tracing experiments. We generated a triple mutant mouse model, by crossing EF1α-Gsα^R201C^ mice with *BAF53b-Cre* mice and *R26-mTmG mice*, to obtain *BAF53b-GFP;EF1α-Gsα^R201C^* mice (Fig. 3A). In this model, which developed a FD skeletal phenotype completely overlapping that of EF1α-Gsα^R201C^ mice^29^, all nerve fibers in the body could be traced based on the expression of GFP. Indeed, preliminary analysis of peripheral organs such as eyes, skin, gut and skeletal muscle confirmed the presence of GFP-expressing nerve fibers. Interestingly we did not observe any change in the innervation of these organs between *BAF53b-GFP* and *BAF53b-GFP;EF1α-Gsα^R201C^* mice (Fig. S3A).

**Figure 3.**
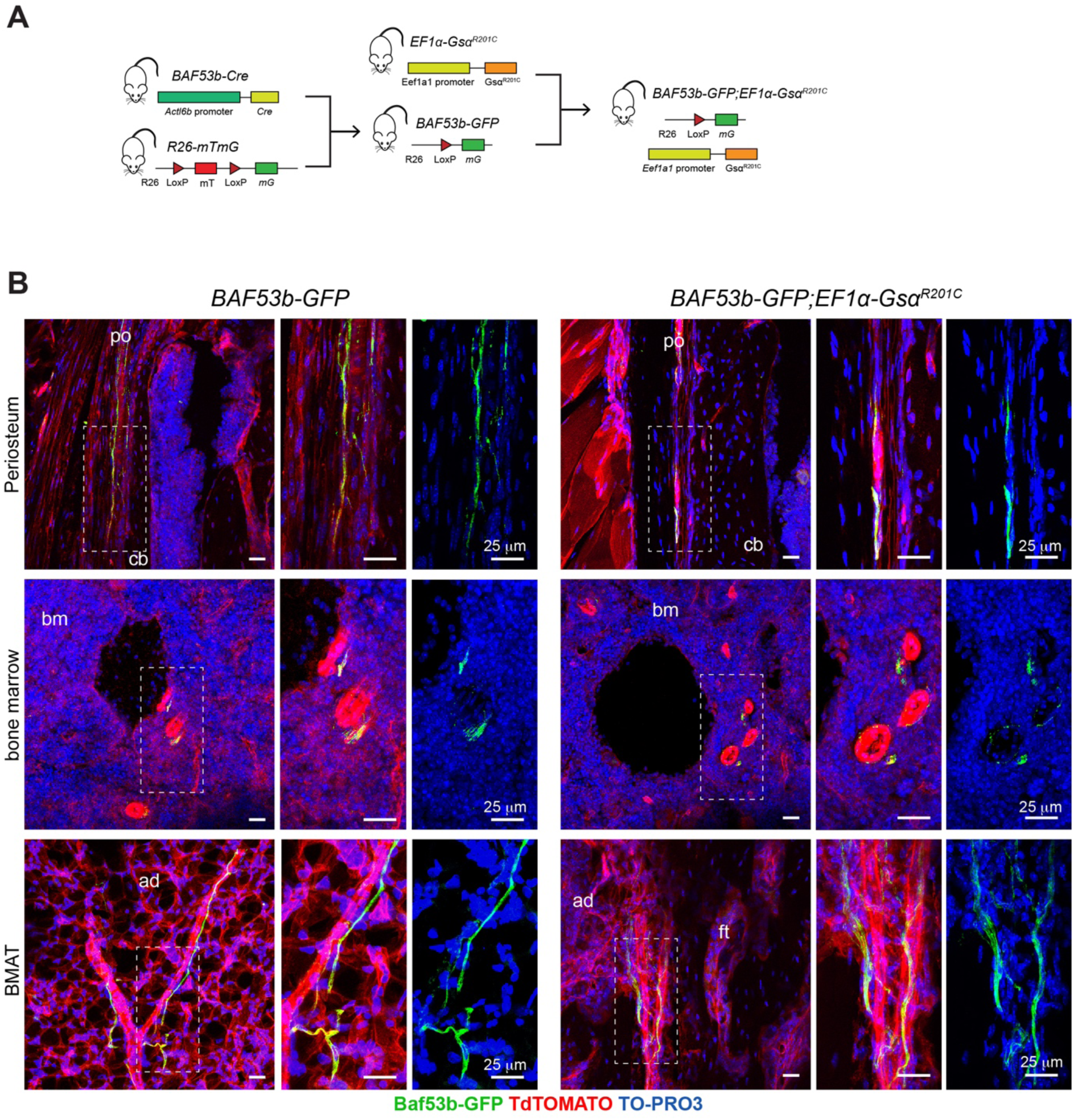
Localization of GFP+ nerve fibers in the periosteum and bone marrow of 8-month-old *BAF53b-GFP* (controls) and *BAF53b-GFP;EF1α-Gsα^R201C^* mice. A) Generation of the triple transgenic *BAF53b-GFP;EF1α-Gsα^R201C^* mouse model of FD. B) Nerve fibers distribution in the fibrous layer of the periosteum, hematopoietic bone marrow and in the bone marrow adipose tissue (BMAT) of caudal vertebrae. No overall differences between BAF53b-GFP and *BAF53b-GFP;EF1α-Gsα^R201C^* mice are observed. Femoral periosteum and BMAT panels are from longitudinal sections, femoral hematopoietic bone marrow images are from transverse sections. Z-stacks of the confocal images are 60 μm thick. po=periosteum, cb=cortical bone, bm=bone marrow, ad=bone marrow adipocytes, ft=fibrous tissue.

In femurs and caudal vertebrae of *BAF53-GFP* control mice, nerve fibers were found mainly in the fibrous layer of the periosteum (Fig. 3B, 4A). GFP positive fibers were also detected in the hematopoietic bone marrow of femurs and in the fatty bone marrow of caudal vertebrae (Fig. 3B) In contrast, they were only rarely found in the cortical bone of the same skeletal segments (Fig. S3B). As expected, nerve fibers were never observed in the growth plate (Fig. S3C). In *BAF53b-GFP;EF1α-Gsα^R201C^* mice, the pattern of bone innervation in skeletal compartment not involved by FD, was overall similar to that of control mice, with periosteum being the most innervated compartment (Fig. 3B). We never observed sprouting of nerve fibers.

The distribution of nerve fibers within the FD tissue was analyzed in lesions of different skeletal segments, such as femurs, calvariae, jaws and caudal vertebrae. Overall, we did not observe a massive innervation of FD lesions (Fig. 4A-C, S3C, D). Indeed, GFP-positive nerve fibers were focally detected in the fibrotic marrow (Fig. 4A-C), showing mainly a linear morphology (Fig. 4B, C) and a regular distribution along the blood vessel wall (Fig. 4B, C and 5A). According, images reminiscent of nerve fiber sprouting were observed rarely and only at sites of blood vessel branching (Fig. 4C iii and vi). We never observed neuroma-like structures.

**Figure 4.**
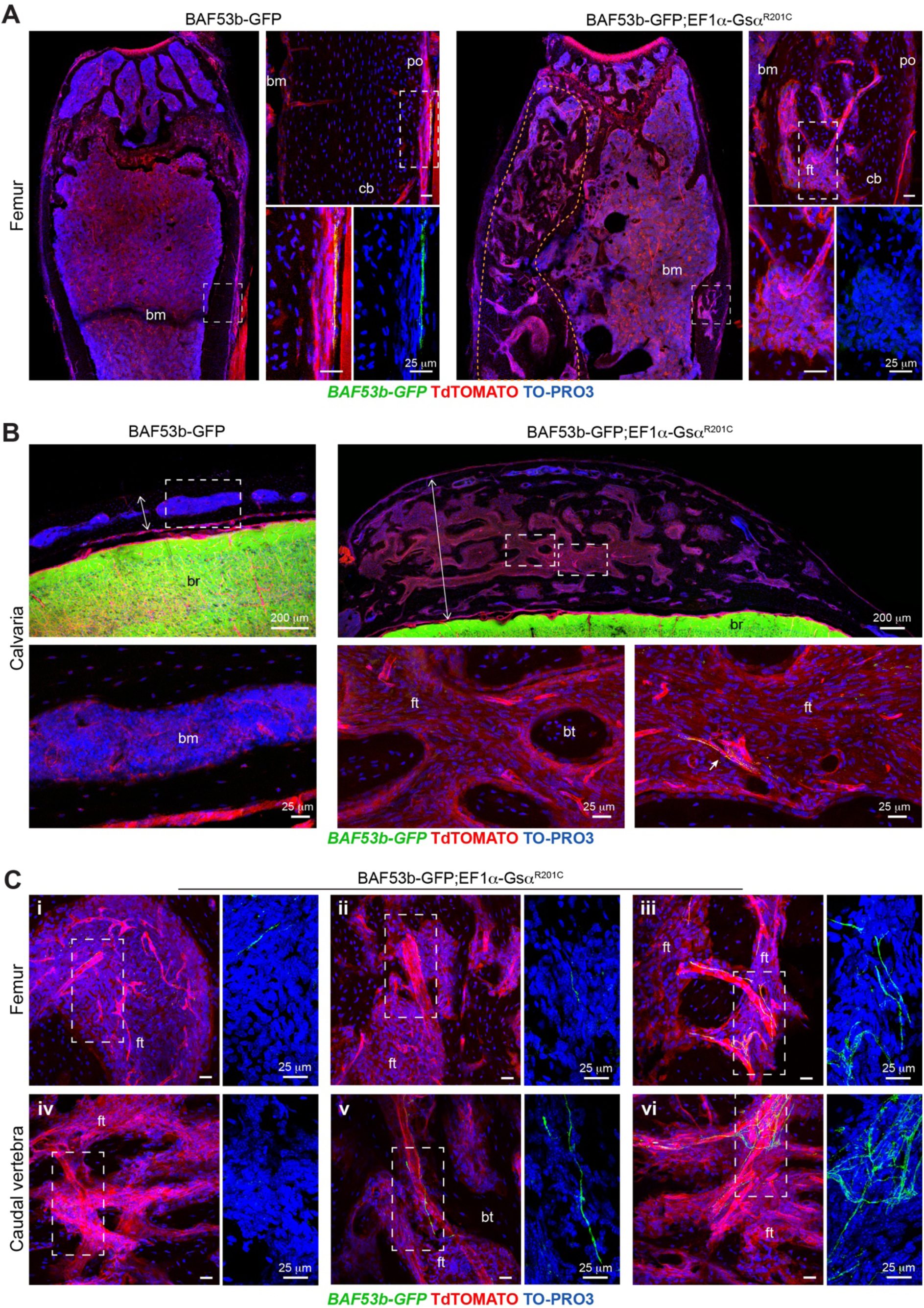
Localization of GFP+ nerve fibers in mouse FD lesions. A) Low and high magnification images of femurs, showing the lack of an evident innervation in these FD lesions of *Baf53b-GFP;Ef1α-Gsα^R201C^* mice. FD lesion in the femur of *Baf53b-GFP;Ef1α-Gsα^R201C^* mouse is delimited by the orange dotted line. B) Low and high magnification images of calvariae, showing a drastic morphological change of the bone microarchitecture due to the development of FD in *Baf53b-GFP;Ef1α-Gsα^R201C^* mice, and the presence of rare nerve fibers within the fibrous tissue (arrow). Note the markedly increased thickness of the calvarial bone in *Baf53b-GFP;Ef1α-Gsα^R201C^* mouse. C) Representative high magnification images taken in the metaphyseal regions of femurs (i-iii) and throughout the caudal vertebra (iv-vi) showing the high heterogeneity in the presence of GFP+ nerve fibers in the fibrotic marrow of *Baf53b-GFP;Ef1α-Gsα^R201C^* mice. Please note the intimate and necessary association with large caliber blood vessels. Z-stacks of the confocal images are 60 μm thick. po=periosteum, cb=cortical bone, bm=bone marrow, ft=fibrous tissue, bt=bone trabecula.

SCA-1 and CD31 immunolocalization showed that GFP-expressing nerve fibers were restricted to the wall of arterial blood vessels (Fig. 5A, B). Quantification of nerve and blood vessel profiles further supported the indispensable perivascular distribution of nerve fibers in the fibrous tissue since all GFP+ axons were found to be associated to arteries/arterioles (Fig. 5C). Interestingly, the fraction of the vessel wall surface coated by nerve fibers was lower in FD lesions of *BAF53b-GFP;EF1α-Gsα^R201C^* mice compared to bone marrow of control mice (Fig. 5D) indicating that the florid vascularization of FD tissue is not necessarily associated with the expansion of the peripheral nerve network.

**Figure 5.**
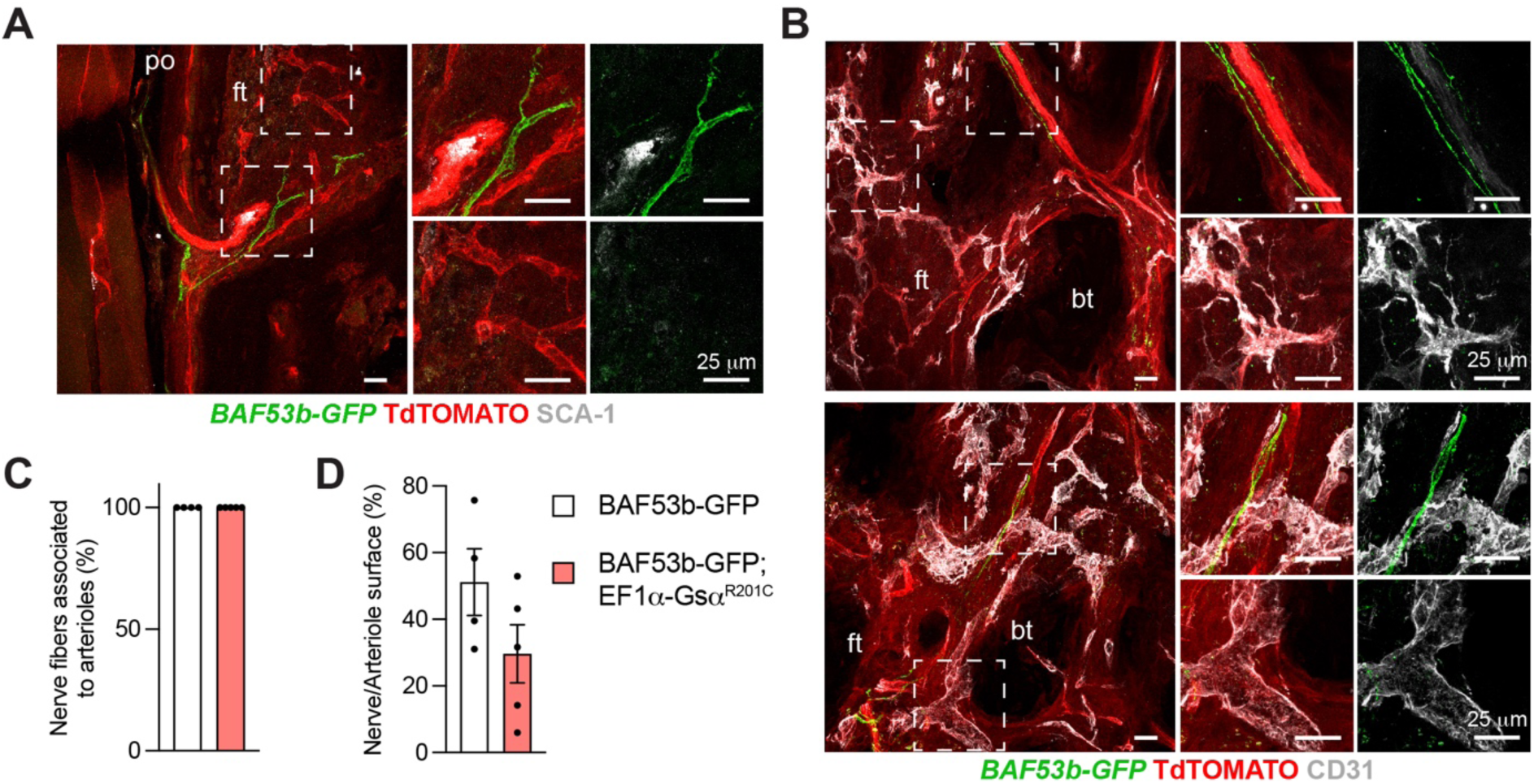
Association of nerve fibers with arterial blood vessels in mouse FD lesions. A, B) Representative confocal images of FD lesions from caudal vertebrae of 8-month-old *Baf53b-GFP;Ef1α-Gsα^R201C^* mice immunostained with SCA-1 (A) and CD31 (B) antibodies. GFP+ nerve fibers are predominantly associated with SCA-1+ arteries/arterioles within the fibrotic marrow. C) Quantification of nerve fibers associated to arteriolar surface. Please note that all the nerve fibers in *Baf53b-GFP* and *Baf53b-GFP;Ef1α-Gsα^R201C^* mice were found next to arterial blood vessels. D) Percentage of nerve fiber surface on arteriolar surface. Note the lower ratio in *Baf53b-GFP;Ef1α-Gsα^R201C^* mice compared to *Baf53b-GFP* control mice, indicating that neoangiogenesis in FD lesions is not necessarily accompanied by nerve fiber growth. po=periosteum, ft=fibrous tissue, bt=bone trabecula.

### Sensory nerve fibers are found within FD bone lesions

Since GFP labeling in *Baf53b-GFP;Ef1α-Gsα^R201C^* mice did not allow to distinguish the different types of nerve fibers, the presence of sensory innervation was assessed by immunolocalization studies based on the expression of NF200, CGRP and TRKA.

We observed that the pattern of immunoreactivity was overall comparable in periosteum, cortical bone and bone marrow of WT and EF1α-Gsα^R201C^ mice (Fig. 6A, S4A, B). Importantly, NF200 as well as CGRP and TRKA were found to be expressed by the nerve fibers observed within the fibrous tissue of EF1α-Gsα^R201C^ mice (Fig 6B), demonstrating the presence of sensory innervation within FD lesions.

**Figure 6.**
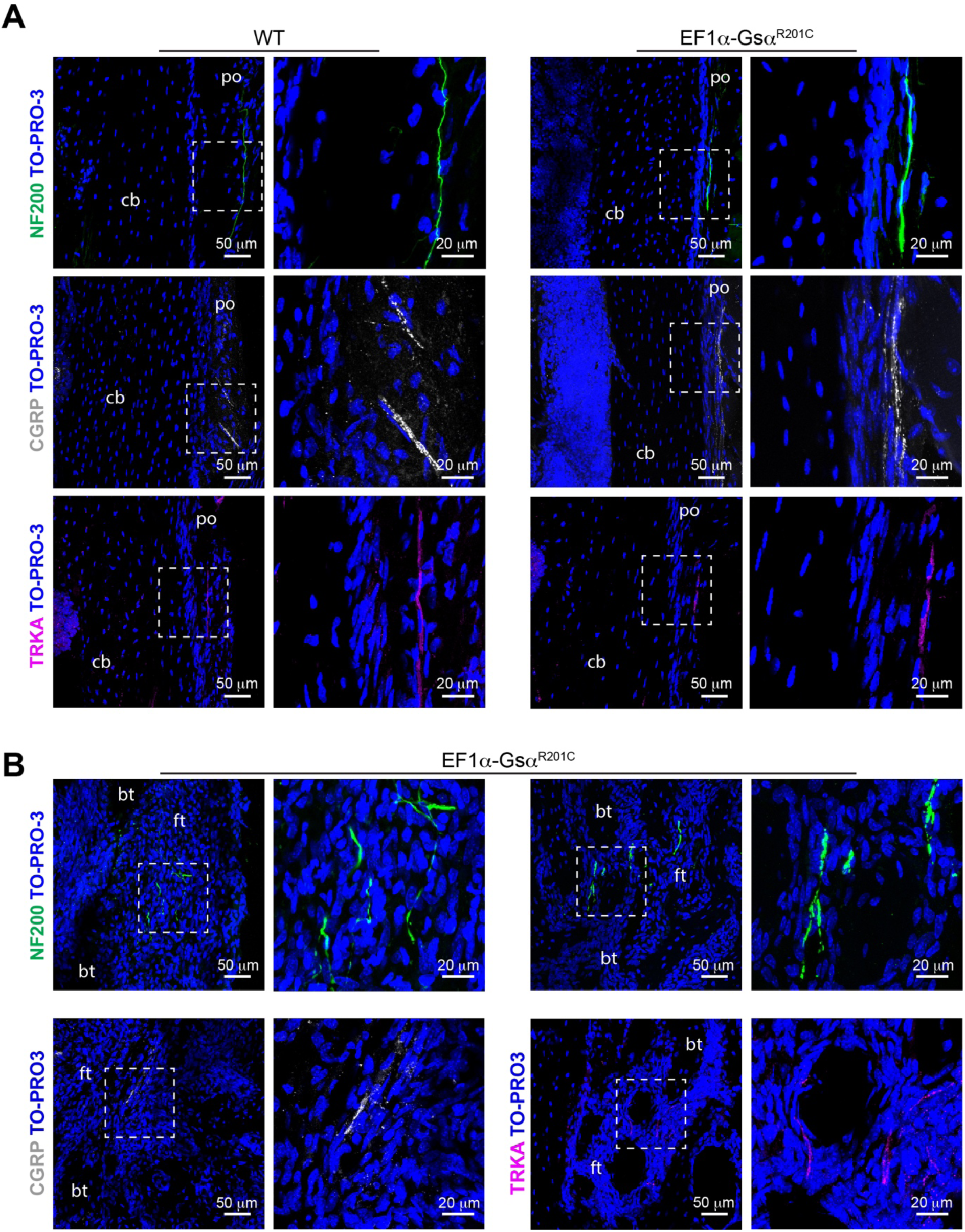
Sensory nerve fibers in mouse FD lesions. A) Confocal images of tibial periosteum of WT littermates and EF1α-Gsα^R201C^ mice immunostained with NF200, CGRP and TRKA antibodies showing similar representation of the different nerve markers in the two groups of mice. B) Representative confocal images of tibial FD lesions of EF1α-Gsα^R201C^ mice showing NF200+, CGRP+ and TRKA+ nerve fibers within the pathological fibrous tissue. po=periosteum, cb=cortical bone, ft=fibrous tissue, bt=bone trabecula.

### Sensory nerve fibers are rarely found within human FD bone lesions

We then investigated the innervation pattern of 13 bone biopsies obtained from different skeletal sites of 7 patients diagnosed with FD/MAS. The biopsy area ranged from 9.73 to 291.12 mm^2^ with the largest representing a whole-mount transverse section of a tibia (Fig. 7A). All samples showed the typical fibro-osseous tissue replacing normal bone and marrow (Fig 7A-D and S5). Surprisingly, immunolocalization of NF200 on paraffin embedded sections did not show extensive labeling within the pathological tissue in our series of samples (Fig 7C, S5). We observed NF200 immunoreactivity in intra-lesional nerve trunks associated with large blood vessels near to cortical bone or within the fibrotic marrow (Fig 7D, S5). Rare axons within the same structures also expressed CGRP (Fig. 7D). As with NF200, no CGRP immunoreactivity was detected in the rich network of blood vessels expanding within the fibrous tissue (Fig. 7C). To improve the sensitivity of the analysis, immunolocalization of NF200 was performed on thick gelatin-embedded sections followed by confocal Z-stack analysis, which revealed only rare and thin nerve fibers within the fibrotic marrow (Fig. 7E). These experiments demonstrate that FD lesions, at least from young patients, may be devoid of nerve fibers, suggesting that nerve sprouting or formation of neuroma-like structures is not a pathological feature of the disease.

**Figure 7.**
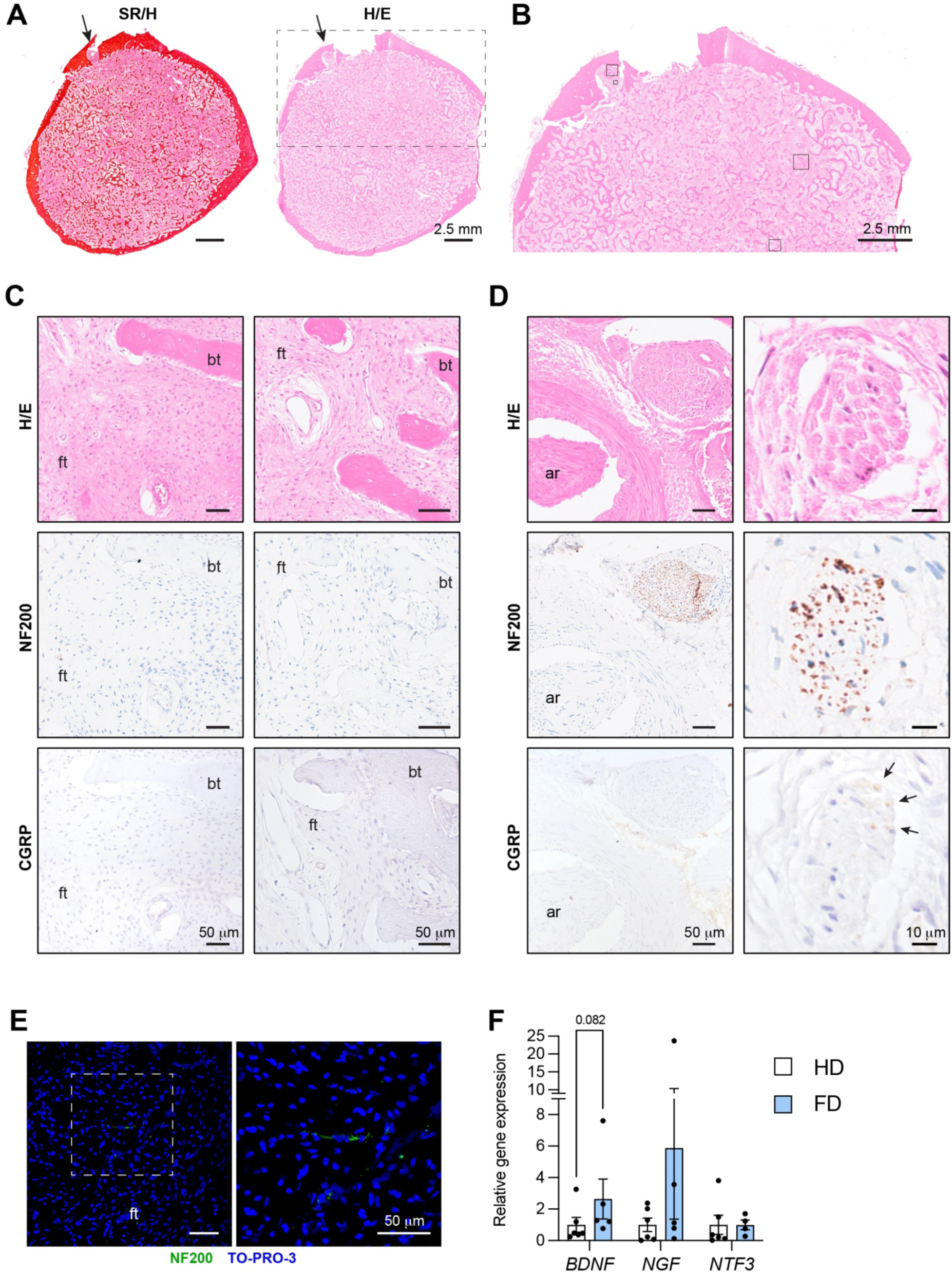
Pattern of innervation in human FD lesions. A) Representative sirius red/hematoxylin (SR/H) and hematoxylin/eosin (H/E)-stained transverse section of a human tibia with FD. A nerve structure was identified within the fibrous tissue, close to the cortical bone (arrow). The dotted box is a reference for the image in B. B) High magnification image of the H/E-stained section. Boxes are references for the images in C and D. C) Images from 3 μm-thick paraffin sections stained with H/E and immunostained with NF200 and CGRP showing the absence of nerve fibers within the fibrous tissue. D) Images from 3 μm-thick paraffin sections stained with H/E and immunostained with NF200 and CGRP showing peripheral nerves identified at the endosteal area of the sample, in proximity of a large artery (ar). Please not that while NF200 staining was found in all the nerve fibers identified, CGRP staining was found only in some axons of the fascicle (arrows). E) Representative images from 60 μm-thick gelatin sections immunostained with NF200, showing very rare, thin nerve fibers in the fibrous tissue. F) qPCR gene expression analysis of neurotrophins performed on fresh healthy donors (HD) and FD bone tissues. *RNA18SN5* was used as housekeeping gene for normalization. Data are shown as dot plots with column bars showing all the experimental samples. Statistical analysis was performed using Student t-test and exact p-value for *BDNF* panels is reported. ft=fibrous tissue, bt=bone trabecula, ar=artery.

Since we previously observed that the levels of *BDNF*, a neurotrophin involved in pain mechanisms, is increased in FD tissue compared to normal bone, we expanded the analysis to other neurothrophins. Confirming our previous results *BDNF* transcripts were higher in FD than in HD bone samples, although the result didn’t reach the statistical significance due to the high variability^17,26^. Similarly, *NGF* expression was higher in FD compared to HD, although variability was observed amongst the different FD samples. No differences were observed in *NTF3* expression, while *NTF4* expression was undetectable in both FD and HD bone samples (Fig. 7F).

## Discussion

Pain is defined as “an unpleasant sensory and emotional experience associated with, or resembling that associated with, actual or potential tissue damage” (https://www.iasp-pain.org/). Chronic bone pain is a clinical feature of many genetic skeletal diseases but in most of them the driving mechanisms remain unknown. This gap of knowledge has multiple reasons, including the technical hurdle in studying bone and bone marrow innervation and the difficulty in gathering large cohorts of patients with rare bone diseases.

FD is a genetic disorder in which bone pain is often caused by bone deformity and fracture, as in appendicular bones^30–32^ or nerve impingement, as in craniofacial bones^33^. However, even in the absence of these complications, many FD patients complain of an “intrinsic”, chronic pain^8,9,11,34,35^, which in a subset of patients is of neuropathic type^9^. Interestingly, the skeletal disease burden, which correlates with the patient’s quality of life and illness perception^34^, is predictive for the development of pain^35^ but does not affect the pain score^8,9,36^. Therefore, understanding the physio-pathological bases of the unpleasant sensory experience associated with this disease is particularly intricate.

The EF1α-Gsα^R201C^ mice^29^ is currently the mouse model that more closely reproduces the natural history of the human bone disease^29^. In FD patients, bone lesions are not detected at birth^37^ and appear at variable times and sites during skeletal growth^33,38^. Similarly, in EF1α-Gsα^R201C^ mice bone lesions develop spontaneously and asynchronously in the post-natal life and generate different patterns of skeletal involvement. Therefore, these mice offer a suitable model for correlation studies between disease burden and other clinical or biochemical parameters. Furthermore, due to the ubiquitous expression of the transgene, they allow to investigate the cell-autonomous effects of the Gsα mutation on any cell type and compartment, including the peripheral and central nervous systems.

In this study, we first analyzed EF1α-Gsα^R201C^ mice using the burrowing and nesting tests and in both of them we observed a spontaneous pain-like behavior, which developed early in adult mice (3 months) and was fully expressed by 5 months of age. This behavior did not depend on the number and distribution of FD lesions and, at least in the burrowing test, appeared earlier in female compared to male mice. These results are consistent with the notion that the severity of the skeletal phenotype is not the main pain-determinant factor in FD^9,39,40^ and that the pain experience is overall more pronounced in female than male patients^9^. Of note, burrowing activity and nest construction in mice are comparable to human functions of daily living including care of the self and interaction with domestic and social environment^22^. Therefore, the impaired burrowing and nesting capacity of EF1α-Gsα^R201C^ mice seems to reflect the overall physical impairment that may be caused by FD^41^. Similarly, the tail flick scores were in complete agreement with data from human FD patients. Indeed, a recent work by Golden at al. reported a higher pain tolerance to heat and cold stimuli in FD/MAS patients compared to controls^10^. Of note, the authors of this study properly raised the point of the origin of tolerance to noxious stimuli in FD patients, asking whether it might result from the continuous use of analgesics or from an adaptation process to persistent pain. The analysis of our transgenic mice, which did not receive any analgesic treatment, seems to support the latter possibility.

We then investigated, for the first time, the pattern of bone innervation in FD. To this aim, on the one side we developed a new EF1α-Gsα^R201C^ mouse model expressing a reporter gene under the control of a pan-neuronal promoter; on the other side, we analyzed the innervation status of a series of human FD lesions. Lineage tracing studies clearly demonstrated that the overall amount and distribution of nerve fibers in organs and bones of mice with mutated Gsα were comparable to those of controls.

Within mouse FD bone lesions, sensory innervation consisted essentially in linear nerve fibers that were associated with arterial blood vessels. All human FD biopsies showed very few intra-lesional fibers, despite the rich vascularization of the fibrotic tissue and regardless of the levels of expression of neurotropic factors. Altogether these data seem to dispel the hypothesis that the Gsα mutation and/or the growth of the FD tissue stimulate sprouting of sensory fibers as observed in bone cancer^2,42–44^. Nonetheless, the presence of sensory innervation within the fibrous tissue indicates the possibility of signal transmission to CNS. It is interesting to note that the variable and unpredictable representation of nerve fibers within different lesions is consistent with the lack of correlation between pain and disease burden that characterizes the disease. Undoubtedly, however, additional mechanisms are required to generate the unpleasant sensation experienced by FD patients. Previous work showed that the anatomical distribution, rather than the number and size of FD lesions, is an important determinant in the clinical expression of the disease. Indeed, painful lesions are more frequently observed in lower extremities and ribs compared to upper extremities and craniofacial bones^35^. The relevance of lesion topography could depend on the size or density of pre-existing nerves in affected skeletal segments^1^ as in thoracic lesions, and/or the presence of mechanical stress, as in lesions of lower extremities^35^. However, it must be noted that very different pain scores were reported in patients with craniofacial FD and comparable images of trigeminal compression and displacement^36^. Similarly, we did not observe a clear connection between mouse behavioral scores and the anatomical distribution of FD. Specific pathological changes and/or inflammatory cytokine secretion within individual FD lesions may also be implicated in the generation of an algogenic micro-environment. Inflammatory cytokines such as IL-6 have been shown to be produced by the fibrous tissue^45^. However, IL-6 is not elevated in the circulation^46^, and neither IL-6 inhibition nor other anti-inflammatory drugs, decrease significantly bone pain in FD patients^11,47^. Importantly, inflammatory cells are never found within the FD lesion microenvironment, except for mastocytes^48^. Enhanced bone resorption, which is a recurrent feature of FD, may also play a role by generating an acidic environment that stimulates sensory neurons^49,50^. Since the number of osteoclasts varies among different FD lesions^17,26^ it is likely that lesions with active bone resorption result in higher levels of acidity and enhanced nerve stimulation compared to quiescent ones. However, pain-relief is not achieved in all FD patients treated with anti-bone resorption drugs^12–14,51,52^. Furthermore, osteomalacia may contribute to the pain sensation in FD, as inhibition of FGF23 using burosumab was accompanied by pain reduction in a child and adult patients^53,54^. Finally, considering the somatic condition of the human disease, the presence of unpleasant sensation reported by some patients might depend on the expression of the Gsα mutation in critical components of the nervous system. Our analyses of FD mouse skeleton and organs, seem to exclude a cell-autonomous effect of the mutation on the growth and structure of peripheral nerves. However, it remains to assess whether the mutation alters the function of sensory nerves and/or it affects the organization and activity of the central nervous system, where pain-related information is elaborated. Interestingly, Golden et al recently analyzed a cohort of FD patients using diffusor tension imaging and observed a structural compromission of some withe matter pathways that are critical to personality and other feelings such as depression and anxiety^10^. Although discordant reports are found in literature about the role of these feelings in FD, the work by Golden and colleagues suggests that the sensation of chronic pain in FD patients may result from the complex integration of physical, psychological and neurobiological mechanisms. This would be consistent with a nociplastic type of pain, in which altered nociception is not explained by tissue damage or pathological alteration of the sensory system (https://www.iasp-pain.org/)^55^. Although it is obviously difficult to explore the role played by psychological states and negative feelings caused by self or societal stigma in mice, interesting information have emerged from our work. Specifically, the results of the open field test strongly suggested that anxiety was not increased in mice with FD lesions compared to controls.

In conclusion, we showed that chronic bone pain and physical impairment are intrinsic and reproducible features of FD. Furthermore, we demonstrated for the first time that, albeit sensory fibers are found within FD lesions and may contribute to bone pain, abnormal nerve sprouting and formation of neuroma-like structures are not pathological features of this disease. This is a preliminary study that has evident limitations, including the absence of data on FD mouse central nervous system, the low number of human FD biopsies and the absence of rigorous scale-based evaluation of pain in FD patients from which they were obtained. Nonetheless, it provides novel findings that will help to better characterize the unpleasant sensory experience associated with FD.

## Acknowledgments

The authors are thankful to Gino Gaudio, passed away in December 2023, for his help in building the tubes for the burrowing test.

## References

1. Steverink, J. G. et al. Sensory Innervation of Human Bone: An Immunohistochemical Study to Further Understand Bone Pain. J Pain 22, 1385–1395 (2021).

2. Luger, N. M., Mach, D. B., Sevcik, M. A. & Mantyh, P. W. Bone Cancer Pain: From Model to Mechanism to Therapy. J Pain Symptom Manage 29, 32–46 (2005).

3. Chartier, S. R., Mitchell, S. A. T., Majuta, L. A. & Mantyh, P. W. The Changing Sensory and Sympathetic Innervation of the Young, Adult and Aging Mouse Femur. Neuroscience 387, 178–190 (2018).

4. Castañeda-Corral, G. et al. The majority of myelinated and unmyelinated sensory nerve fibers that innervate bone express the tropomyosin receptor kinase A. Neuroscience 178, 196–207 (2011).

5. Brazill, J. M., Beeve, A. T., Craft, C. S., Ivanusic, J. J. & Scheller, E. L. Nerves in Bone: Evolving Concepts in Pain and Anabolism. Journal of Bone and Mineral Research 34, 1393–1406 (2019).

6. Collins, M. T., Boyce, A. M. & Riminucci, M. *Primer on the Metabolic Bone Diseases and Disorders of Mineral Metabolism*. In Bilezikian JP (ed) Primer of Metabolic bone diseases 9th ed (Wiley, 2018). doi:10.1002/9781119266594.

7. Boyce, A. M. & Collins, M. T. Fibrous Dysplasia/McCune-Albright Syndrome: A Rare, Mosaic Disease of Gαs Activation. Endocr Rev 41, 345–370 (2020).

8. Kelly, M. H., Brillante, B. & Collins, M. T. Pain in fibrous dysplasia of bone: age-related changes and the anatomical distribution of skeletal lesions. Osteoporosis International 19, 57–63 (2008).

9. Spencer, T. L. et al. Neuropathic-like Pain in Fibrous Dysplasia/McCune-Albright Syndrome. J Clin Endocrinol Metab 107, e2258–e2266 (2022).

10. Golden, E. et al. Phenotyping Pain in Patients With Fibrous Dysplasia/McCune-Albright Syndrome. J Clin Endocrinol Metab 109, 771–782 (2024).

11. Chapurlat, R. D. et al. Pathophysiology and medical treatment of pain in fibrous dysplasia of bone. Orphanet J Rare Dis 7, S3 (2012).

12. Boyce, A. M. et al. Denosumab treatment for fibrous dysplasia. J Bone Miner Res 27, 1462–70 (2012).

13. Majoor, B. C. J. et al. Denosumab in Patients With Fibrous Dysplasia Previously Treated With Bisphosphonates. J Clin Endocrinol Metab 104, 6069–6078 (2019).

14. Meier, M. E. et al. Safety of therapy with and withdrawal from denosumab in fibrous dysplasia and McCune-Albright syndrome: an observational study. Journal of Bone and Mineral Research 36, 1729–1738 (2021).

15. Corsi, A. et al. Zoledronic Acid in a Mouse Model of Human Fibrous Dysplasia: Ineffectiveness on Tissue Pathology, Formation of ‘Giant Osteoclasts’ and Pathogenetic Implications. Calcif Tissue Int 107, 603–610 (2020).

16. Palmisano, B. et al. RANKL Inhibition in Fibrous Dysplasia of Bone: A Preclinical Study in a Mouse Model of the Human Disease. Journal of Bone and Mineral Research 34, 2171–2182 (2019).

17. Palmisano, B. et al. A pathogenic role for brain-derived neurotrophic factor (BDNF) in fibrous dysplasia of bone. Bone 181, 117047 (2024).

18. Collins, M. T. et al. An Instrument to Measure Skeletal Burden and Predict Functional Outcome in Fibrous Dysplasia of Bone. Journal of Bone and Mineral Research 20, 219–226 (2005).

19. Zhan, X. et al. Generation of <scp> BAF </scp> 53b-<scp> C </scp> re transgenic mice with pan-neuronal <scp>C</scp> re activities. genesis 53, 440–448 (2015).

20. Muzumdar, M. D., Tasic, B., Miyamichi, K., Li, L. & Luo, L. A global double-fluorescent Cre reporter mouse. genesis 45, 593–605 (2007).

21. Roedel, A., Storch, C., Holsboer, F. & Ohl, F. Effects of light or dark phase testing on behavioural and cognitive performance in DBA mice. Lab Anim 40, 371–381 (2006).

22. Deacon, R. Assessing Burrowing, Nest Construction, and Hoarding in Mice. Journal of Visualized Experiments (2012) doi:10.3791/2607.

23. Deacon, R. M. Assessing nest building in mice. Nat Protoc 1, 1117–1119 (2006).

24. Palmisano, B. et al. GsαR201C and estrogen reveal different subsets of bone marrow adiponectin expressing osteogenic cells. Bone Res 10, 50 (2022).

25. Schindelin, J., et al. Fiji: an open-source platform for biological-image analysis. Nat Methods 9, 676–682 (2012).

26. Persichetti, A. et al. Nanostring technology on Fibrous Dysplasia bone biopsies. A pilot study suggesting different histology-related molecular profiles. Bone Rep 16, 101156 (2022).

27. Seibenhener, M. L. & Wooten, M. C. Use of the Open Field Maze to Measure Locomotor and Anxiety-like Behavior in Mice. Journal of Visualized Experiments (2015) doi:10.3791/52434.

28. Carter, M. & Shieh, J. Animal Behavior. in Guide to Research Techniques in Neuroscience 39–71 (Elsevier, 2015). doi:10.1016/B978-0-12-800511-8.00002-2.

29. Saggio, I. et al. Constitutive expression of Gsα(R201C) in mice produces a heritable, direct replica of human fibrous dysplasia bone pathology and demonstrates its natural history. J Bone Miner Res 29, 2357–68 (2014).

30. Nakashima, Y., Kotoura, Y., Nagashima, T., Yamamuro, T. & Hamashima, Y. Monostotic fibrous dysplasia in the femoral neck. A clinicopathologic study. Clin Orthop Relat Res 242–8 (1984).

31. Leet, A. I. et al. Fracture Incidence in Polyostotic Fibrous Dysplasia and the McCune-Albright Syndrome. Journal of Bone and Mineral Research 19, 571–577 (2004).

32. Benhamou, J., Gensburger, D., Messiaen, C. & Chapurlat, R. Prognostic Factors From an Epidemiologic Evaluation of Fibrous Dysplasia of Bone in a Modern Cohort: The FRANCEDYS Study. Journal of Bone and Mineral Research 31, 2167–2172 (2016).

33. Collins, M. T. Spectrum and natural history of fibrous dysplasia of bone. J Bone Miner Res 21 **Suppl 2**, P99–P104 (2006).

34. Majoor, B. C. J. et al. Illness Perceptions are Associated with Quality of Life in Patients with Fibrous Dysplasia. Calcif Tissue Int 102, 23–31 (2018).

35. Majoor, B. C. J. et al. Pain in fibrous dysplasia: relationship with anatomical and clinical features. Acta Orthop 90, 401–405 (2019).

36. Golden, E., et al. Case Report: The Imperfect Association Between Craniofacial Lesion Burden and Pain in Fibrous Dysplasia. Front Neurol 13, (2022).

37. Corsi, A., et al. Neonatal McCune-Albright Syndrome: A Unique Syndromic Profile With an Unfavorable Outcome. JBMR Plus 3, (2019).

38. Robinson, C., Collins, M. T. & Boyce, A. M. Fibrous Dysplasia/McCune-Albright Syndrome: Clinical and Translational Perspectives. Curr Osteoporos Rep 14, 178–186 (2016).

39. Guerin Lemaire, H., Merle, B., Borel, O., Gensburger, D. & Chapurlat, R. Serum periostin levels and severity of fibrous dysplasia of bone. Bone 121, 68–71 (2019).

40. Florenzano, P. et al. Age-Related Changes and Effects of Bisphosphonates on Bone Turnover and Disease Progression in Fibrous Dysplasia of Bone. Journal of Bone and Mineral Research 34, 653–660 (2019).

41. Kelly, M. H., Brillante, B., Kushner, H., Gehron Robey, P. & Collins, M. T. Physical function is impaired but quality of life preserved in patients with fibrous dysplasia of bone. Bone 37, 388–394 (2005).

42. Mantyh, W. G. et al. Blockade of nerve sprouting and neuroma formation markedly attenuates the development of late stage cancer pain. Neuroscience 171, 588–598 (2010).

43. Bloom, A. P. et al. Breast Cancer-Induced Bone Remodeling, Skeletal Pain, and Sprouting of Sensory Nerve Fibers. J Pain 12, 698–711 (2011).

44. Diaz-delCastillo, M. et al. Metastatic Infiltration of Nervous Tissue and Periosteal Nerve Sprouting in Multiple Myeloma-Induced Bone Pain in Mice and Human. The Journal of Neuroscience 43, 5414–5430 (2023).

45. Riminucci, M. et al. Osteoclastogenesis in fibrous dysplasia of bone: in situ and in vitro analysis of IL-6 expression. Bone 33, 434–442 (2003).

46. Meier, M. E. et al. Clinical value of RANKL, OPG, IL-6 and sclerostin as biomarkers for fibrous dysplasia/McCune-Albright syndrome. Bone 171, 116744 (2023).

47. Chapurlat, R. et al. Inhibition of IL-6 in the treatment of fibrous dysplasia of bone: The randomized double-blind placebo-controlled TOCIDYS trial. Bone 157, 116343 (2022).

48. Bertrand, G., Minard, M. F., Simard, C. & Rebel, A. [Ultrastructural study of a case of monostotic fibrous dysplasia]. Ann Anat Pathol (Paris) 23, 81–9 (1978).

49. Yoneda, T., Hiasa, M., Nagata, Y., Okui, T. & White, F. Contribution of acidic extracellular microenvironment of cancer-colonized bone to bone pain. Biochim Biophys Acta 1848, 2677–84 (2015).

50. Nagae, M. et al. Osteoclasts play a part in pain due to the inflammation adjacent to bone. Bone 39, 1107–1115 (2006).

51. Boyce, A. M. et al. A Randomized, Double Blind, Placebo-Controlled Trial of Alendronate Treatment for Fibrous Dysplasia of Bone. J Clin Endocrinol Metab 99, 4133–4140 (2014).

52. Chapurlat, R. D., Delmas, P. D., Liens, D. & Meunier, P. J. Long-Term Effects of Intravenous Pamidronate in Fibrous Dysplasia of Bone. Journal of Bone and Mineral Research 12, 1746–1752 (1997).

53. Gladding, A., Szymczuk, V., Auble, B. A. & Boyce, A. M. Burosumab treatment for fibrous dysplasia. Bone 150, 116004 (2021).

54. Stelmachowska-Banaś, M., Cylke-Falkowska, K., Zgliczyński, W. & Misiorowski, W. Burosumab treatment for FGF23-related hypophoshatemia in an adult patient with severe fibrous dysplasia in McCune-Albright syndrome. Endocrine Abstracts (2024) doi:10.1530/endoabs.99.P249.

55. Kosek, E. et al. Chronic nociplastic pain affecting the musculoskeletal system: clinical criteria and grading system. Pain 162, 2629–2634 (2021).

