## Supplementary files for "Bone pain in Fibrous dysplasia does not rely on aberrant sensory nerve sprouting or neuroma formation"

**Supplementary data**

**Figure S1**

**A**

| Score | Paper used | Description |
|---|---|---|
| 1 | <10% | The shredded paper is not noticeably touched. |
| 2 | <50% | Less than 50% of the shredded paper is used to build a nest |
| 3 | 50-90% | More than 50% of the shredded paper is used to build a nest but several paper strips remain scattered around the cage and the material may be in a broadly defined nest area |
| 4 | >90% | More than 90% of the shredded paper is used to build a nest but the nest is flat and walls not higher than mouse height |
| 5 | >90% | More than 90% of the shredded paper is used to build a nest and the nest is well-organized with walls higher than mouse height |

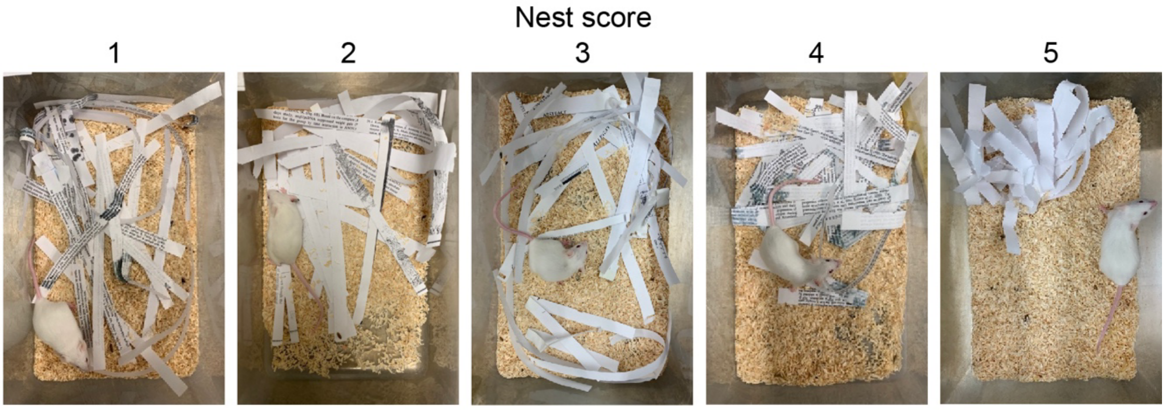


**B**

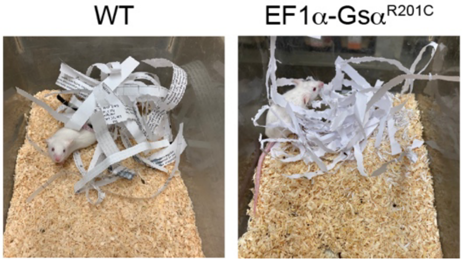


**Figure S1. Nesting test scoring system**. A) Table with scoring and example pictures of the nests. B) Representative high scored nests from WT and EF1$\alpha$-Gs$\alpha^{R201C}$ mice. Note that the shredded paper used for the nest by the EF1$\alpha$-Gs$\alpha^{R201C}$ mouse resulted chewed.

**Figure S2**

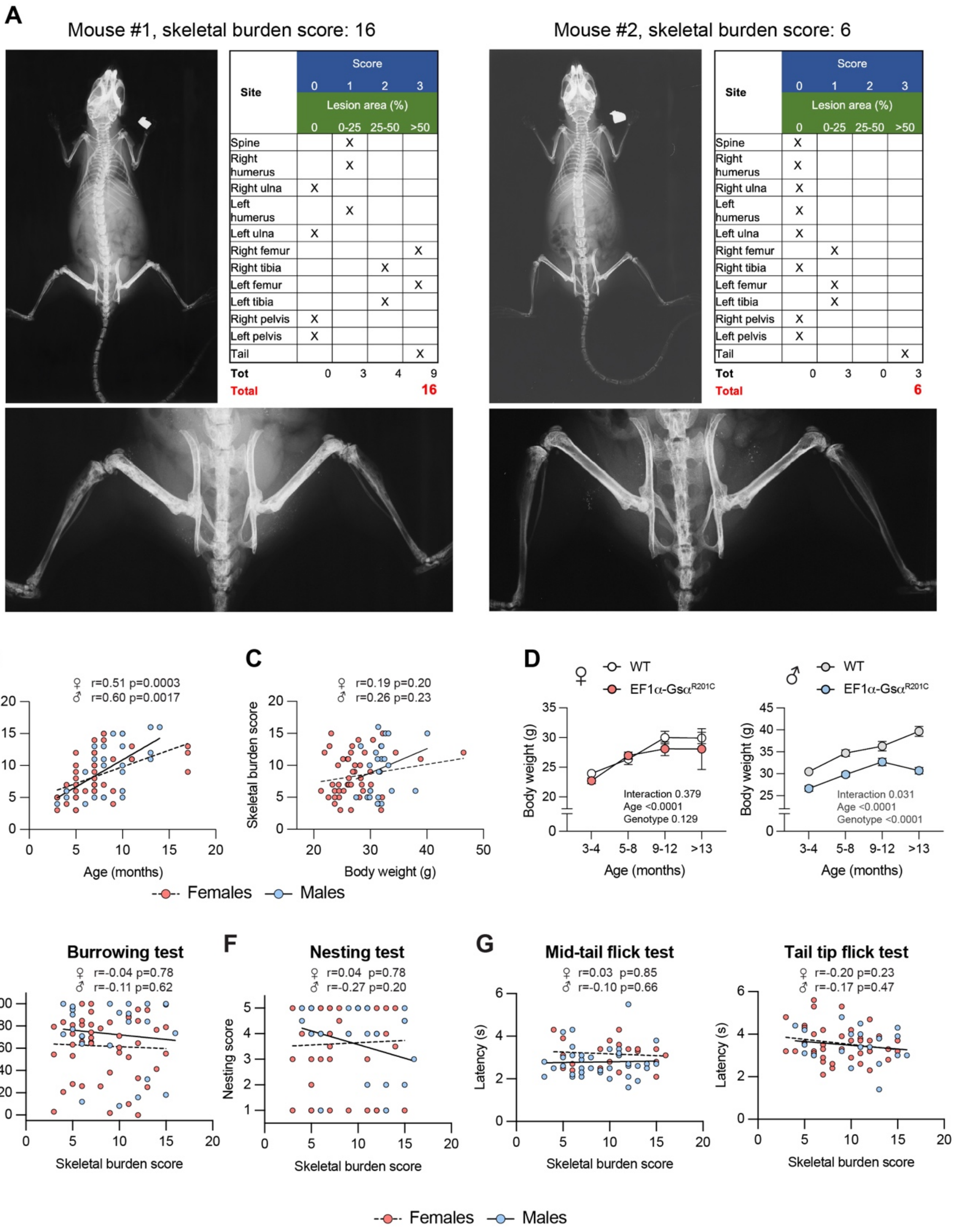


Mouse #1:

| Site | Score 0 / Lesion area 0 | Score 1 / Lesion area 0-25 | Score 2 / Lesion area 25-50 | Score 3 / Lesion area >50 |
|---|---|---|---|---|
| Spine | | X | | |
| Right humerus | | X | | |
| Right ulna | X | | | |
| Left humerus | | X | | |
| Left ulna | X | | | |
| Right femur | | | | X |
| Right tibia | | | X | |
| Left femur | | | | X |
| Left tibia | | | X | |
| Right pelvis | X | | | |
| Left pelvis | X | | | |
| Tail | | | | X |
| Tot | 0 | 3 | 4 | 9 |
| Total | | | | 16 |

Mouse #2:

| Site | Score 0 / Lesion area 0 | Score 1 / Lesion area 0-25 | Score 2 / Lesion area 25-50 | Score 3 / Lesion area >50 |
|---|---|---|---|---|
| Spine | X | | | |
| Right humerus | X | | | |
| Right ulna | X | | | |
| Left humerus | X | | | |
| Left ulna | X | | | |
| Right femur | | X | | |
| Right tibia | X | | | |
| Left femur | | X | | |
| Left tibia | | X | | |
| Right pelvis | X | | | |
| Left pelvis | X | | | |
| Tail | | | | X |
| Tot | 0 | 3 | 0 | 3 |
| Total | | | | 6 |

**Figure S2. Skeletal burden score evaluation and correlation with pain behavior and other parameters in EF1α-Gsα$^{R201C}$ mice**. A) Radiograms and table for evaluation of the skeletal disease burden. In these pictures, two representative mice are shown with different scores. B) Correlation analysis between burden scores and age. C) Correlation analysis between burden scores and body weight. D) Measurements of body weight in female and male WT and EF1α-Gsα$^{R201C}$ mice at different ages. P-values from the two-way ANOVA analysis are reported in each graph. E) Correlation analysis between burrowing capacity and skeletal burden score. F) Correlation analysis between nesting and burden scores. G) Correlation analysis between tail flick latency times and burden scores. In B, C and E-G correlation coefficient (r) and p-value are shown above each graph. Please note that mice with very low disease burden scores can show poor burrowing and nesting ability; similarly, mice with very high disease burden scores may not experience a pain-like behavior.

Figure S3

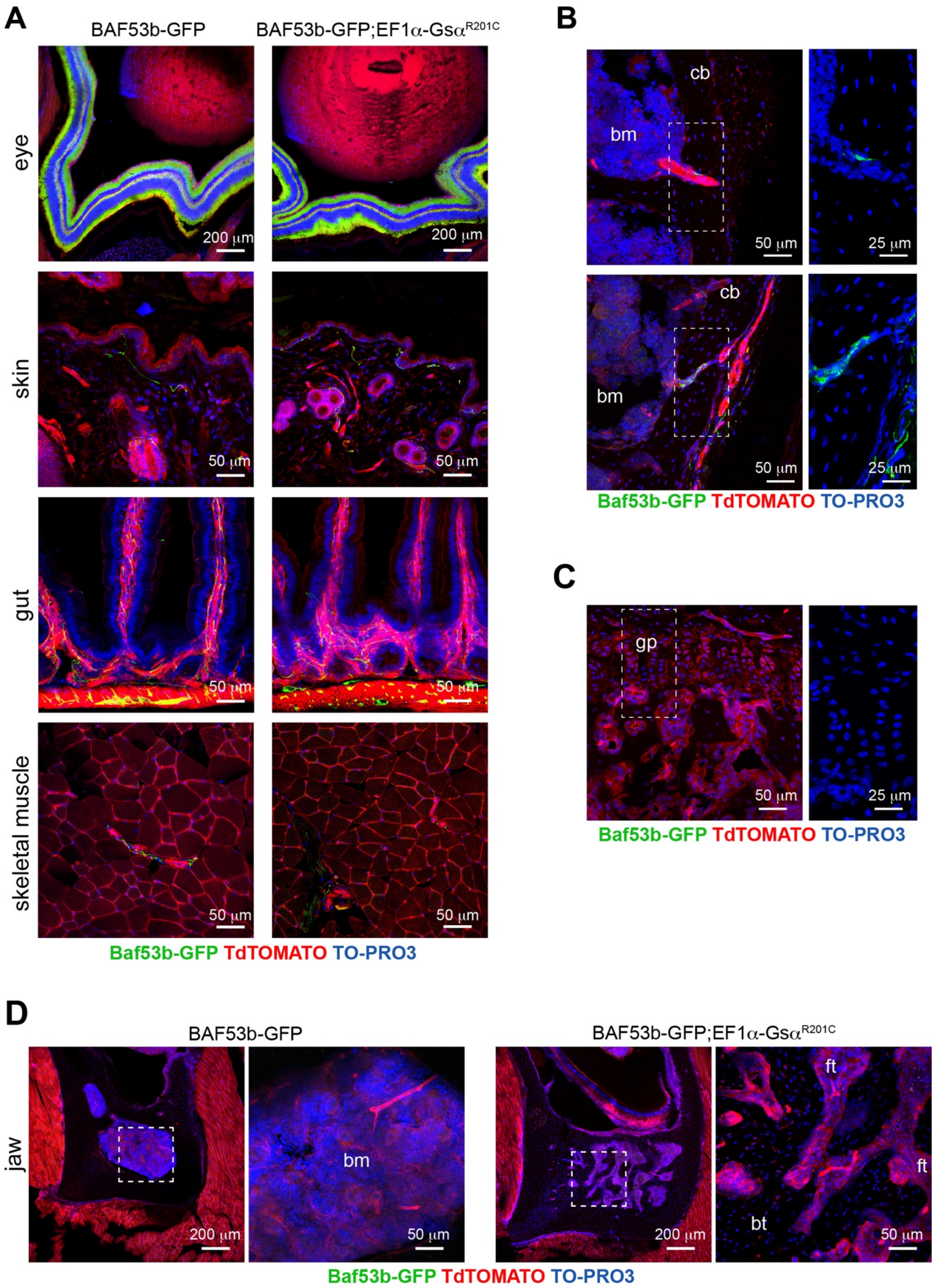


**Figure S3. Pattern of Baf53b-GFP tracing in different organs and skeletal compartments.** A) GFP labeling in some peripheral organs, revealing no overall differences between *BAF53b-GFP* and *BAF53b-GFP;EF1α-Gsα*$^{R201C}$ mice. B) Representative pictures showing GFP+ intracortical nerve fibers in *BAF53b-GFP* mice. C) Representative pictures showing the absence of nerve fibers in the cartilaginous growth plate and trabecular bone. D) Pictures of jaw bones showing no nerve fibers neither in the hematopoietic bone marrow of *BAF53b-GFP* mice nor in fibro-osseous lesions fo *BAF53b-GFP;EF1α-Gsα*$^{R201C}$ mice. Except for the muscle images showing a single plane, Z-stacks of the confocal images are 60 μm thick. cb=cortical bone, bm=bone marrow, ft=fibrous tissue, bt=bone trabecula, gp=growth plate.

**Figure S4**

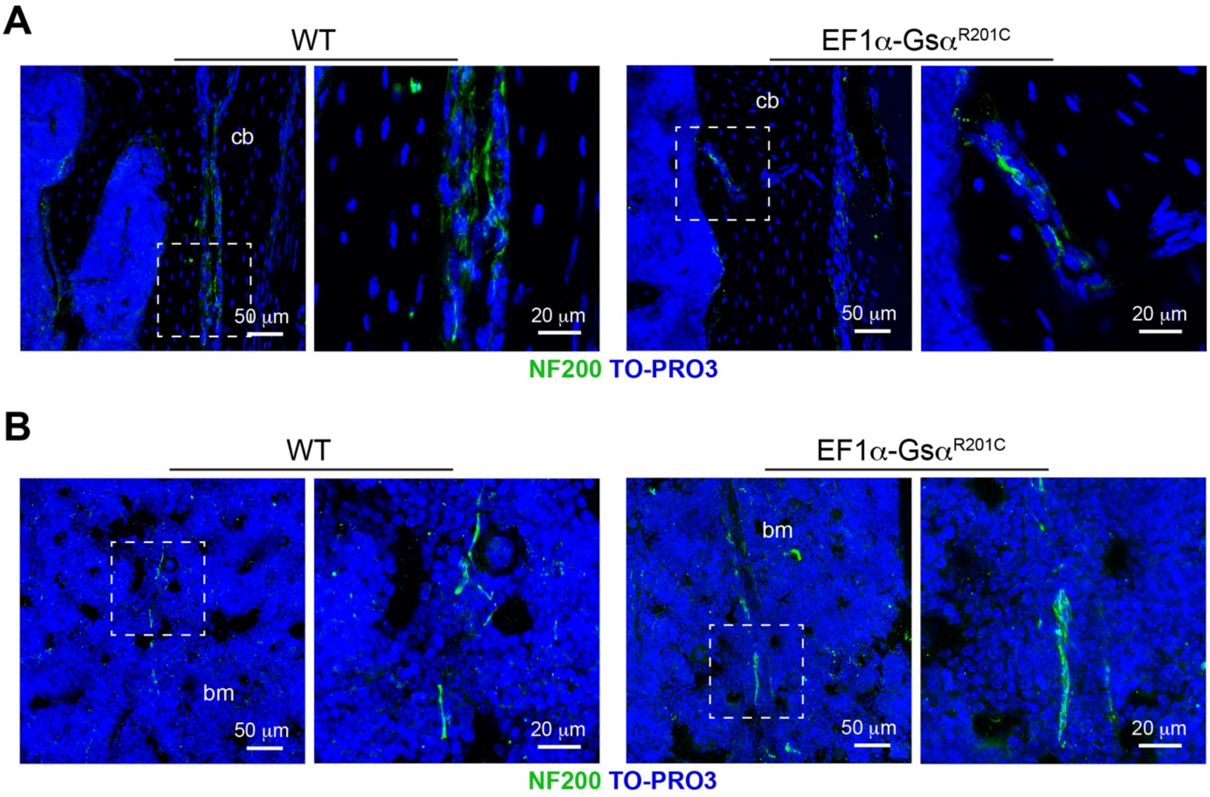


**Figure S4. Similar pattern of sensory innervation in cortical bone and bone marrow in WT and EF1α-Gsα$^{R201C}$ mice.** A) Representative NF200 immunostaining of tibial cortical bone. B) Representative NF200 immunostaining of hematopoietic bone marrow showing linear nerve fibers in both genotypes. Z-stacks of the confocal images are 60 μm thick. cb=cortical bone, bm=bone marrow.

**Figure S5.**

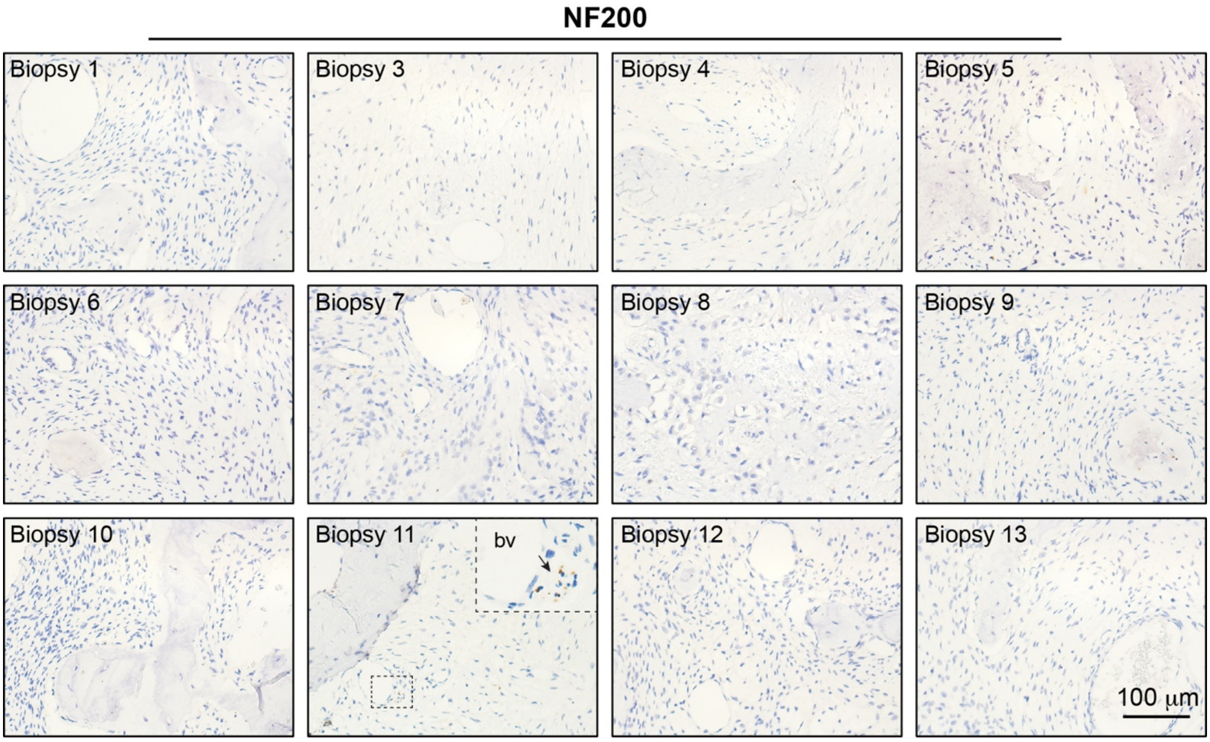


**Figure S5.** Representative images of all the human FD bone biopsies immunostained with NF200 antibody, showing the lack of sensory nerve fiber sprouting within the pathological FD tissue. A small nerve trunk is seen in Biopsy 11 (arrow). bv=blood vessel. Please note that Biopsy 2 is shown in Fig. 7.

**Table S1**

**Mouse Genotyping**

| **Mouse strain** | **Sequence 5' - 3'** |
|---|---|
| *EF1α-Gsα*$^{R201C}$ | F: GGATTACGCGTCCAACAGCG<br>R: TCCCGGATGACCATGTTGTA |
| *Baf53b-Cre* | F: GCATTGCTGTCACTTGGTCGT<br>R: CGATGCAACGAGTGATGAGG |
| *R26-mTmG* | F: CTCTGCTGCCTCCTGGCTTCT<br>R: CGAGGCGGATCACAAGCAATA<br>R: TCAATGGGCGGGGGTCGTT |

**qPCR in human samples**

| **Gene** | **Sequence 5' - 3'** |
|---|---|
| *BDNF* | F: ACGAGACCAAGTGCAATCCC<br>R: TACGACTGGGTAGTTCGGCA |
| *NGF* | F: ACCCGCAACATTACTGTGGACC<br>R: GACCTCGAAGTCCAGATCCTGA |
| *NTF3* | F: CAAGCAGATGGTGGACGTTAAGG<br>R: TCGCAGCAGTTCGGTGTCCATT |
| *NTF4* | F: GCAAGGCTGATAACGCTGAGGA<br>R: CCTGGGCATCAGCGGTCAATG |
| *RNA18SN5* | F: CGATGCTCTTAGCTGAGTGT<br>R: GGTCCAAGAATTTCACCTCT |
